# Glacier-fed stream biofilms harbour diverse resistomes and biosynthetic gene clusters

**DOI:** 10.1101/2021.11.18.469141

**Authors:** Susheel Bhanu Busi, Laura de Nies, Paraskevi Pramateftaki, Massimo Bourquin, Leïla Ezzat, Tyler J. Kohler, Stilianos Fodelianakis, Grégoire Michoud, Hannes Peter, Michail Styllas, Matteo Tolosano, Vincent De Staercke, Martina Schön, Valentina Galata, Paul Wilmes, Tom Battin

## Abstract

**Background:** Antimicrobial resistance (AMR) is a universal phenomenon whose origins lay in natural ecological interactions such as competition within niches, within and between micro- to higher-order organisms. However, the ecological and evolutionary processes shaping AMR need to be better understood in view of better antimicrobial stewardship. Resolving antibiotic biosynthetic pathways, including biosynthetic gene clusters (BGCs), and corresponding antimicrobial resistance genes (ARGs) may therefore help in understanding the inherent mechanisms. However, to study these phenomena, it is crucial to examine the origins of AMR in pristine environments with limited anthropogenic influences. In this context, epilithic biofilms residing in glacier-fed streams (GFSs) are an excellent model system to study diverse, intra- and inter-domain, ecological crosstalk.

**Results:** We assessed the resistomes of epilithic biofilms from GFSs across the Southern Alps (New Zealand) and the Caucasus (Russia) and observed that both bacteria and eukaryotes encoded twenty-nine distinct AMR categories. Of these, beta-lactam, aminoglycoside, and multidrug resistance were both abundant and taxonomically distributed in most of the bacterial and eukaryotic phyla. AMR-encoding phyla included Bacteroidota and Proteobacteria among the bacteria, alongside Ochrophyta (algae) among the eukaryotes. Additionally, BGCs involved in the production of antibacterial compounds were identified across all phyla in the epilithic biofilms. Furthermore, we found that several bacterial genera (*Flavobacterium*, *Polaromonas*, etc.) including representatives of the superphylum Patescibacteria encode both ARGs and BGCs within close proximity of each other, thereby demonstrating their capacity to simultaneously influence and compete within the microbial community.

**Conclusions:** Our findings highlight the presence and abundance of AMR in epilithic biofilms within GFSs. Additionally, we identify their role in the complex intra- and inter-domain competition and the underlying mechanisms influencing microbial survival in GFS epilithic biofilms. We demonstrate that eukaryotes may serve as AMR reservoirs owing to their potential for encoding ARGs. We also find that the taxonomic affiliation of the AMR and the BGCs are congruent. Importantly, our findings allow for understanding how naturally occurring BGCs and AMR contribute to the epilithic biofilms mode of life in GFSs. Importantly, these observations may be generalizable and potentially extended to other environments which may be more or less impacted by human activity.

## Background

Today, antimicrobial resistance (AMR) has become a well-known threat to human health with an estimated number of 700,000 people per year dying of drug-resistant infections [1]. The dramatic rise of antimicrobial resistance over the past decade has even led to the moniker, “silent pandemic” [2]. Therefore, AMR is often directly associated with human impacted environments with a global increase in resistant bacteria linked to the over- and mis-use of antibiotics [3]. However, contrary to public perception, AMR is a natural phenomenon, which has existed for billions of years [4]. Long before the rather recent use of antibiotics in the clinical setting, microorganisms have used these, along with corresponding protective mechanisms, to establish competitive advantages over other microbes contending for the same environment and/or resources [5].

Microbes, in general, produce a range of secondary metabolites with diverse chemical structures which in turn confer a variety of functions, including antibiotics [6]. Such secondary metabolites including metal transporters and quorum sensing molecules [7,8] are not directly associated with the growth of microorganisms themselves but instead are known to provide benefits by acting as growth inhibitors against competing bacteria. Consequently, many of these natural products have found their uses in industrial settings as well as in human medicine as anti-infective drugs [7,9,10]. The biosynthetic pathways responsible for producing these specialized metabolites are encoded by locally clustered groups of genes known as ‘biosynthetic gene clusters’ (BGCs). Typically, BGCs include genes for expression control, self-resistance, and metabolite export [11]. They can, however, be further divided into various classes including non-ribosomal peptide synthetases (NRPSs), type I and type II polyketide synthases (PKSs), terpenes, and bacteriocins alongside others [10]. NRPSs and PKSs specifically have been of interest due to their known synthesis of putative antibiotics [12,13]. Furthermore, evidence suggests that within these BGCs at least one resistance gene conferring resistance can be found as a self-defense mechanism against the potentially harmful secondary metabolites encoded by the BGC [14]. For instance, the tylosin-biosynthetic gene cluster of *Streptomyces fradiae* also encodes three resistance genes (*tlrB*, *tlrC* and *tlrD*) [15], while in another example, *Streptomyces toyacaensis*, the *vanHAX* resistance cassette is proximal to the vancomycin biosynthesis gene cluster, thereby encoding inherent resistance [16].

Remote and pristine microbial communities provide a rich genetic resource to explore the historical evolutionary origins of naturally occurring antibiotic resistance from the pre-antibiotic era. Only in few pristine environments with limited anthropogenic influence (e.g., permafrost, glaciers, deep sea, and polar regions) can remnants of the above-described ancient biological warfare mechanisms still be detected. These ARGs and resistant bacteria evolving in pristine environments may therefore be considered the inherent antibiotic resistance present in the environment [5].

We have recently reported the genomic and metabolic adaptations of epilithic biofilms to windows of opportunities in glacier-fed streams (GFSs) [17]. For example, given the short flow season during glacial melt, i.e. summer, the incentive to reproduce quickly while conditions are favourable, is high. During these windows of opportunity, the necessity for taxa to not only acquire physical niches, but also appropriate resources yields a competitive environment. Within these biofilms, we observe complex cross-domain interactions between microorganisms to potentially mitigate the harsh nutrient and environmental conditions of the GFSs. Additionally, owing to their complex biodiversity [18] and generally oligotrophic conditions [19], epilithic biofilms are ideal model systems for understanding BGCs and AMR. While oligotrophy may provide the basis for competition over resources amongst microorganisms such as prokaryotes and (micro-)eukaryotes. Our previous insights revealed that taxa such as *Polaromonas*, *Acidobacteria*, and *Methylotenera* have strong interactions with eukaryotes such as algae and fungi [17]. The inherent diversity allows for understanding the influence of AMR in microbial interactions. For example, the accidental discovery of penicillin by Alexander Fleming in 1928 based on bacterial-fungal interactions, [20], has since been expanded upon by Netzker *et al*. [21]. They reported that microbial interactions lead to the production of bioactive compounds including antibiotics that may shape the microbial consortia within a community.

Here, to shed light on the role of AMR in shaping microbial communities within (relatively) pristine environments, we used high-resolution metagenomics to investigate twenty-one epilithic biofilms from glacier-fed streams. These samples were collected from 8 GFSs spread across the Southern Alps in New Zealand and the Caucasus in Russia (Supplementary Table 1). Herein, we found 29 categories of ARGs within the GFSs across both bacterial and eukaryotic domains. Importantly, most of the AMR was found in bacteria. We also identified antibacterial BGCs that were encoded both in bacterial and eukaryotes suggesting extensive intra- and inter-domain competition. Our findings demonstrate that microorganisms within biofilms from pristine environments not only encode ARGs, but that they may potentially influence several features of epilithic biofilms such as biofilm formation, community assembly and/or maintenance, including conferring mechanisms for competitive advantages under extreme conditions.

## Methods

### Sampling and biomolecular extractions

Eight GFSs were sampled in early- to mid-2019 from the New Zealand Southern Alps and the Russian Caucasus, respectively, for a total of 21 epilithic biofilms (Supp. Table 1). The biofilm samples were collected from each stream reach due to biofilms ranging from abundant to absent, depending on stream geomorphology. One to three biofilm samples were collected per reach (Supp. Table 1), taken using sterilized metal spatulas to scrape rocks, followed by their immediate transfer to cryovials. Samples were immediately flash-frozen in liquid nitrogen and stored at −80 °C until DNA was extracted. DNA from the epilithic biofilms was extracted using a previously established protocol [22] adapted to a smaller scale due to relatively high DNA concentrations. DNA quantification was performed for all samples with the Qubit dsDNA HS kit (Invitrogen).

### Sequencing and data processing for metagenomics

Random shotgun sequencing was performed on all epilithic biofilm DNA samples after library preparation using the NEBNext Ultra II FS library kit. 50 ng of DNA was enzymatically fragmented for 12.5 min and libraries were prepared with six PCR amplification cycles. An average insert of 450 bp was maintained for all libraries. Qubit was used to quantify the libraries followed by sequencing at the Functional Genomics Centre Zurich on a NovaSeq (Illumina) using a S4 flowcell. The metagenomic data was processed using the Integrated Meta-omic Pipeline (IMP v3.0; commit# 9672c874 available at https://git-r3lab.uni.lu/IMP/imp3) [23]. IMP’s workflow includes pre-processing, contig assembly, genome reconstruction (metagenome-assembled genomes, i.e. MAGs) and additional functional analysis of genes based on custom databases in a reproducible manner [23].

### Identification of antimicrobial resistance genes, antibiotic biosynthesis pathways and BGCs

For the prediction of ARGs the IMP-generated contigs were used as input for PathoFact [24]. Identified ARGs were further collapsed into their respective AMR categories in accordance with the Comprehensive Antibiotic Resistance Database (CARD) [25]. PathoFact uses an HMM-based search to identify homologous sequences across genomic data, therefore possibly also detecting resistance genes within eukaryotic genomic fragments. Subsequently, the raw read counts per ORF, obtained from PathoFact, were determined using FeatureCounts [26].

To identify pathways for the biosynthesis of antibiotics, we assigned KEGG orthology (KOs) identifiers to the ORFs using a hidden Markov model [27] (HMM) approach using *hmmsearch* from HMMER 3.1 [28] with a minimum bit score of 40. Additionally, we linked the identified KOs to their corresponding KEGG orthology pathways and extracted the pathways annotated as antibiotic biosynthesis pathways by KEGG. Both the identified ARGs and KEGG pathways were then further linked to associated bacterial taxonomies. The bacterial and eukaryotic taxonomies were assigned using the PhyloDB and MMETSP databases associated with EUKulele (commit# fb8726a; available at https://github.com/AlexanderLabWHOI/EUKulele). Consensus taxonomy per contig was then used for downstream analyses including association with ARGs.

We further identified BGCs within the MAGs using antiSMASH (ANTIbiotics & Secondary Metabolite Analysis SHell) [29] and annotated these using deepBGC [30]. To link BGCs and ARGs, we linked the resistance genes to their associated assembled contigs, followed by identifying the corresponding bins (MAGs) to which said contigs belonged.

### Data analysis

The relative abundance of the ORFs was calculated based on the RNum_Gi method described by Hu *et al*. [31]. Figures for the study, including visualizations derived from the taxonomic and functional analyses, were created using version 3.6 of the R statistical software package [32] and using the *tidyverse* package [33]. Alluvial plots were generated using the *ggalluvial* package [34] while heatmaps were generated using the *ComplexHeatmap* package [35] developed for R. The corresponding visualization and analysis code is available at: https://gitr3lab.uni.lu/laura.denies/Rock_Biofilm_AMR.

## Results

### Antimicrobial resistance in a pristine environment

We characterised the resistomes of GFS epilithic biofilms and assessed the distribution of AMR in twenty-one epilithic biofilm samples, across 8 individual glaciers originating from the Southern Alps in New-Zealand (SA1, SA2, SA3 and SA4) and the Caucasus in Russia (CU1, CU2, CU3, CU4). In total, we identified a high number (n=1840) of ARGs within 29 categories of AMR, with similar AMR profiles observed across all GFSs (Fig. 1a, Supp. Fig. 1), except for SA2 and SA3 where the differences were driven by elevated fluoroquinolone, glycopeptide and phenicol resistance, respectively. It is to be noted that while ARGs refer to the genes encoding specific resistance, AMR categories derived from metagenomic data in this context, typically reflect the functional potential associated with respect to the resistance encoded. Of the identified AMR categories, beta-lactam and multidrug resistance (i.e. resistance conferring protection against multiple antibiotic classes), followed by aminoglycoside resistance, were found to be highly abundant in all samples. We subsequently analysed the diversity of ARGs within the various resistance categories and found beta-lactam resistance to represent the largest resistance category, contributing 930 unique ARGs to the resistome. This was followed by multidrug (179 ARGs) and aminoglycoside (176 ARGs) resistance (Supp. Table 2). In contrast, some resistance categories such as polymyxin and pleuromutilin resistance were only detected at very low levels within the epilithic biofilm resistomes.

**Figure 1.**
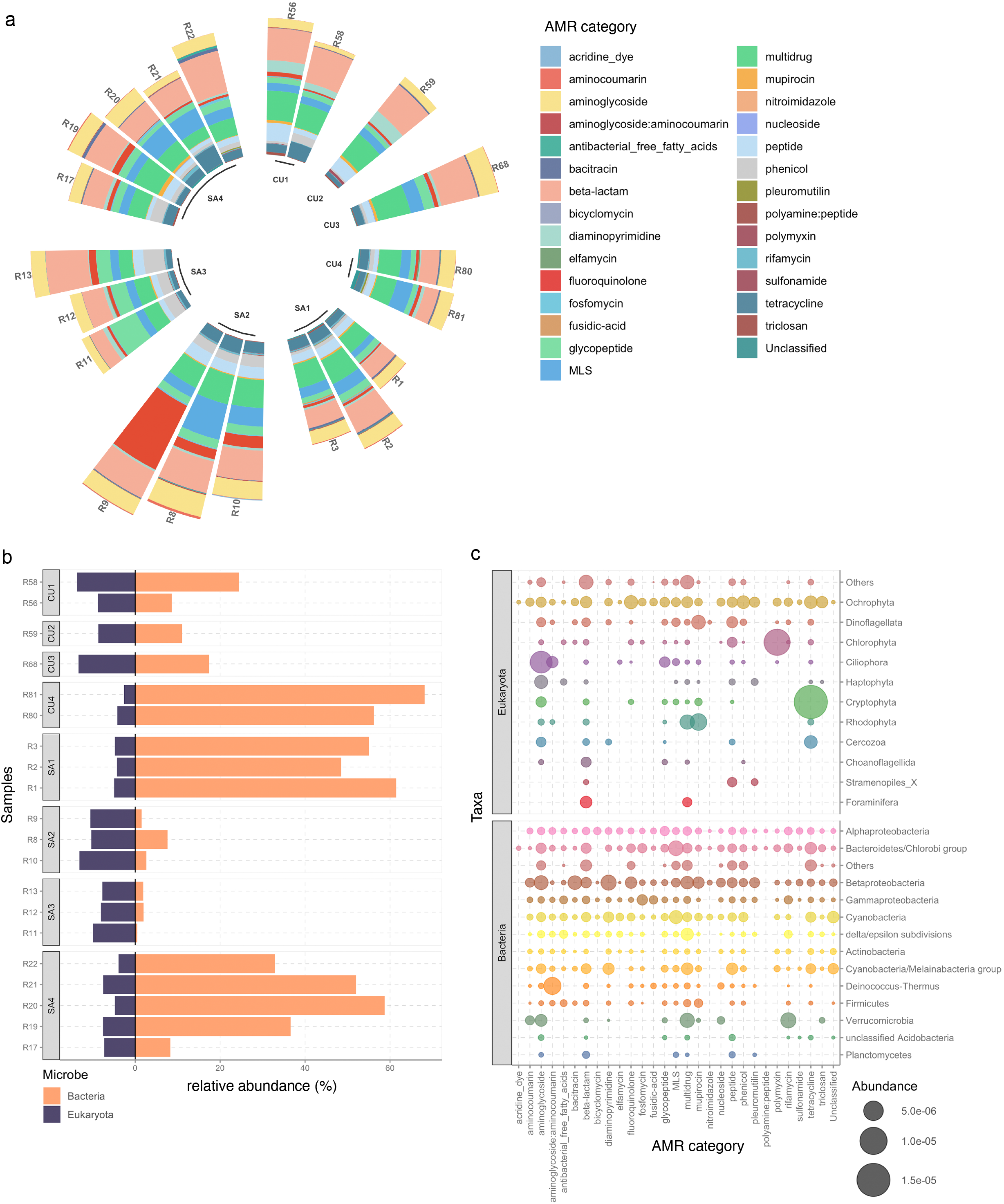
Epilithic biofilms in GFSs harbour a diverse resistome. (a) Relative abundance of 29 AMR categories within 21 epilithic biofilms collected from four New Zealand Southern Alps (SA) and four Russian Caucasus (CU) GFSs. (b) Bar plots depicting the relative abundance of bacteria and eukaryotes encoding ARGs. (c) Phylum-level representation of the AMR abundances across bacteria and eukaryotes. Size of the closed circle indicates the normalised relative abundance (Rnum_Gi; see *Methods*), whereby the color represents individual phyla.

We further investigated the contribution of microbial populations to the resistome and found contributions from both prokaryotes and eukaryotes (Fig. 1b). Prokaryotes within this study refer to bacteria alone, since archaea encoded for an infinitesimal number of ARGs (<0.000001% RNum_GI; *Methods*), and therefore were excluded from further analyses. Among the eukaryotes, the phylum Ochrophyta (algae) was the dominant contributor and encoded most of the AMR categories (Fig. 1c, Supp. Fig. 2a). In bacteria, AMR was more evenly distributed with most of the phyla encoding ARGs across all categories (Fig. 1c). However, members of the Alphaproteobacteria, Betaproteobacteria, and the Bacteroidetes/Chlorobi group encoded the highest overall ARG abundance (Fig. 1c, Supp. Fig. 2b). Additionally, AMR categories such as aminoglycoside, beta-lactam, glycopeptide and rifamycin resistance (among others) were widely distributed in both bacteria as well as among the eukaryotes. On the other hand, categories such as aminocoumarin, bacitracin, and diaminopyrimidine resistance were found to be primarily encoded by bacteria.

### Antibiotic biosynthesis pathways and biosynthetic gene clusters

As described above, beta-lactam, multidrug and aminoglycoside resistance were the most abundant resistance categories within GFS epilithic biofilms. This was not surprising as beta-lactams and aminoglycosides are natural and prevalent compounds [36,37]. Furthermore, multidrug resistance is typically conferred via efflux machineries which were also common in the GFS epilithic biofilms. These typically serve dual purposes in particular for protein export within most bacteria [38]. Based on these results, it is therefore highly likely that pristine environments such as GFSs potentially reflect the spectrum of natural antibiotics and their resistance mechanisms, reinforcing their capacity to serve as natural baselines for assessing enrichments and spread of AMR.

To further understand if these encoded resistance genes reflected natural antibiotic pressure, we investigated pathways associated with antibiotic biosynthesis using the KEGG database [39]. In total, we identified seven different pathways corresponding to the biosynthesis of macrolides (MLS), ansamycins, glycopeptides (vancomycin), beta-lactams (monobactam, penicillin and cephalosporin), aminoglycosides (streptomycin), and tetracyclines, which were present in various abundances in all samples (Supp. Fig. 3a). Importantly, the identified antibiotic synthesis genes thereby corresponded to the resistance categories identified within the epilithic biofilms. Interestingly, in most of the GFSs, antibiotic biosynthesis was primarily encoded by bacteria spanning multiple phyla (Supp. Fig. 3b, Supp. Fig. 3c). Exceptions to these were GL11 and GL15 in which biosynthesis pathways were equally distributed among eukaryotes, specifically Ochrophyta, in addition to bacteria.

To further validate our observations, we assessed the abundance of BGCs, which are known to encode genes for secondary metabolite synthesis, including antibiotics. We found six different structural classes of BGCs by annotating 537 medium-to-high quality (>50% completion and <10% contamination) bacterial and 30 eukaryotic MAGs using antiSmash [29] and DeepBGC [30]. Using this ensemble approach we identified one or more BGCs in most bacterial (n=490, ~91% of all bacterial MAGs) and eukaryotic (n=28) MAGs. Of these BGCs, those annotated with an antibacterial function were dominant across the microbial populations, represented here by the MAGs, and were found across all phyla (Fig. 2a). Overall, a wider variety of BGCs associated with cytotoxic activity, inhibitory, and antifungal mechanisms were also identified in bacteria. Eukaryotes, on the other hand, encoded a high prevalence of antibacterial BGCs (~93% of all eukaryotic MAGs) (Fig. 2a). We further annotated those BGCs identified as antibacterial to determine their subtypes and found that most of them were ‘unknown’ (Fig. 2b). However, other identified subtypes include ribosomally synthesized and post-translationally modified peptides (RiPPs) such as bacteriocins, along with NRPs, PKs, and terpenes.

**Figure 2.**
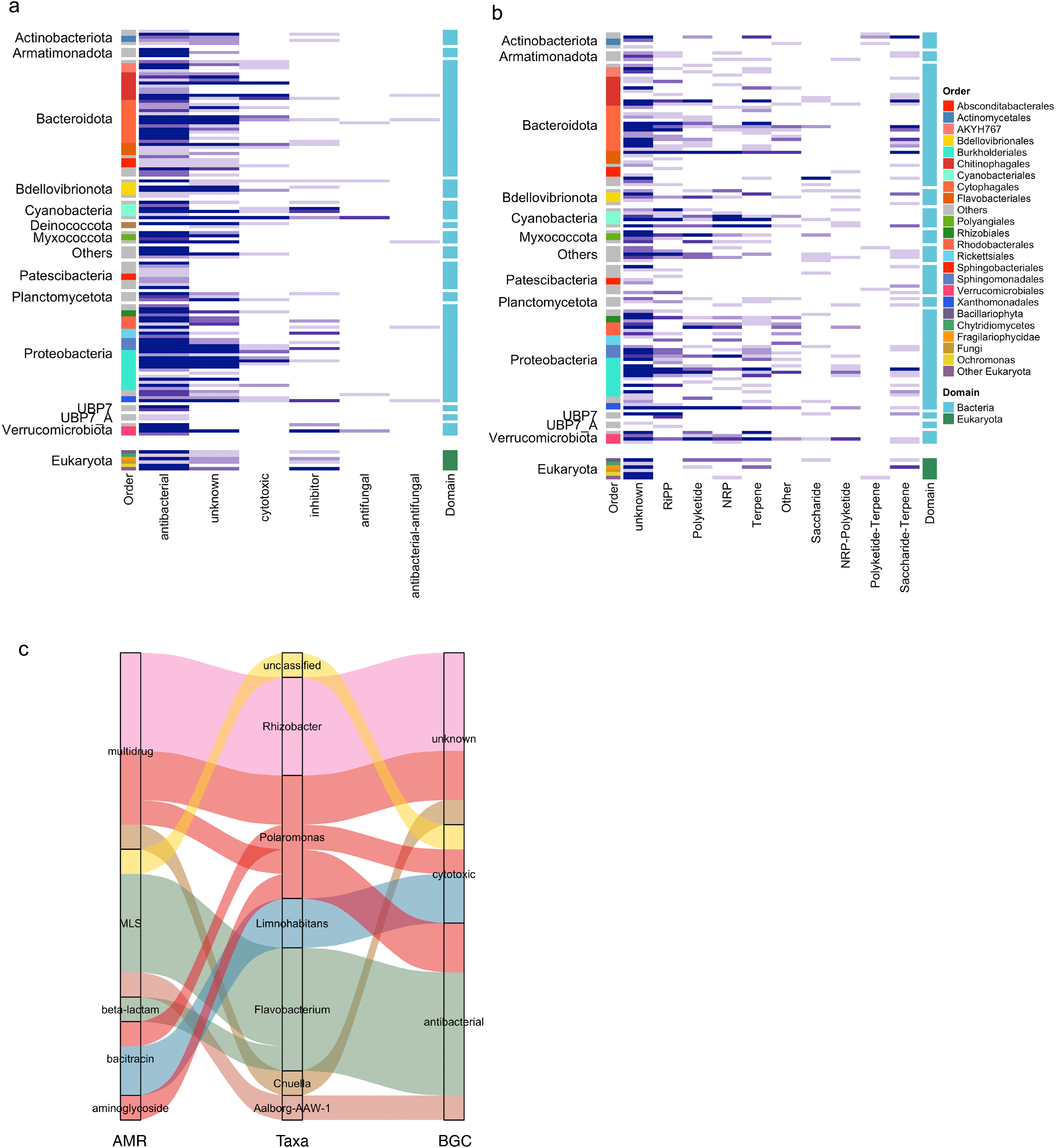
Biosynthetic gene clusters indicate the resistome potential. (a) Heatmap depicting the overall abundance of BGCs identified across bacterial and eukaryotic MAGs. The respective phyla are listed on the left while the coloured legend represents the taxonomic order. (b) In-depth characterisation of the ‘antibacterial’ BGCs found within all phyla and orders across medium-to-high quality MAGs. (c) Alluvial plots depicting the taxa where both BGCs and AMR were found adjacently on the same contig. Colours indicate the genera associated with the MAGs.

According to the resistance hypothesis [14], within or close to, each BGC there is at least one gene conferring resistance to its encoded secondary metabolite. To test this, we assessed whether the MAGs encoding a BGC also encoded corresponding ARGs. In line with this hypothesis, we identified BGCs and their respective resistance genes in close proximity to each other through their localization on the same contig. Consequently, we identified various BGCs encoded together with ARGs in both the bacterial and eukaryotic MAGs. For example, we found that an antibacterial BGC was encoded by *Flavobacterium* spp. on the same contig as both MLS (macrolides, lincosamides and streptogramin) and beta-lactam resistance genes (Fig. 2c). Incidentally, we also found that a candidate phyla radiation (CPR) bacterium (Aalborg-AAW-1; phylum Patescibacteria) also encoded both antibacterial BGC and MLS resistance on the same contig.

## Discussion

Microbial reservoirs in pristine environments, with little to no impact from anthropogenic selection pressures, provide the opportunity to investigate the natural propensity and linked evolutionary origins of AMR. Here, by leveraging high-resolution metagenomics on twenty-one epilithic biofilms, we assessed the resistomes of eight individual GFS epilithic biofilms.

To date, while many studies have looked for novel antibiotics and resistance genes in pristine environments such as the deep sea [40] or the polar regions [41], few have explored the full diversity of antibiotic resistance in such environments [42,43]. Van Goethem *et al*. [44] identified 117 naturally occurring ARGs associated with multidrug, aminoglycoside and beta-lactam resistance in pristine Antarctic soils. Similarly, D’Costa *et al*. [4] identified a collection of ARGs encoding resistance to beta-lactams as well as tetracyclines and glycopeptides in 30,000-year-old Beringian permafrost sediments. In agreement with these previous studies, we identified 29 AMR categories, including the previously mentioned resistance categories, in the studied biofilm communities. Among these, the highest ARG abundance was associated with aminoglycoside and beta-lactam resistance. Our study further suggests that although the overall abundance differs, the epilithic resistome was highly similar in all GFSs, independent of origin (i.e. New Zealand or Russia). Furthermore, our results agree with the results obtained in other resistomes identified in pristine environments such as Antarctic soils and permafrost in terms of the identified ARGs. Unlike previous studies, where ARGs were primarily associated with bacteria, we report for the first time that AMR was associated with both bacteria and eukaryotes in various abundances in environmental samples including GFSs. A previous study by Brown *et al*. [45] reported that the IRS-HR (isoleucyl-tRNA synthetase - high resistance) type gene conferring resistance against mupirocin was identified in *Staphylococcus aureus*. More importantly, they suggested that horizontal gene transfer led to the acquisition of IRS-HR genes by bacteria from eukaryotes [45]. Despite these early reports, the contribution of eukaryotes to most resistomes, including from pristine environments, has largely been unexplored thus far. An exception to this was the report by Fairlamb *et al*. [46] who identified eukaryotic drug resistance, especially encoded by fungi (*Candida* and *Aspergillus*) and parasites (*Plasmodium* and *Trypanosoma*). However, most of these modes of resistance were highly specific towards particular drug treatments [46]. Our results specifically revealed that taxa from the phylum Ochrophyta encoded resistance to 28 AMR categories and this was also reflected in other (micro-)eukaryotes.

Apart from encoded resistance mechanisms, microalgae such as Ochrophyta have been of interest as a source of (new) antimicrobial compounds [47,48]. In line with this, Martins *et al*. suggested that extracts from different microalgae may potentially serve not only as antimicrobial agents, but also as anti-cancer therapeutics. However, our present results suggest that these taxa may also serve as environmental reservoirs for AMR itself. It is however presently unclear whether this phenomenon confers advantages with respect to niche occupation and protection against bacterial infection as well as whether the eukaryotes are sensitive to the antibiotics produced by them.

Studies delving into the origins of AMR have reported that fecal pollution may explain ARG abundances in anthropogenically impacted environments [49]. This phenomenon was also observed by Antelo *et al*. [50] and others [51] who detected ARGs in soils in Antarctica, especially in proximity to scientific bases. Although it is plausible that some of the GFSs sampled in our study may indeed be under anthropogenic influence, in pristine environments, AMR is most likely derived from natural antibiotics produced by microorganisms as a competitive advantage. Microorganisms acquire resistance either as a protective measure against other microorganisms [52,53] or as a self-defense mechanism to prevent inadvertent suicide by damaging metabolites [14]. Accordingly, we found both antibiotic biosynthesis pathways and BGCs within the epilithic resistomes. We identified pathways for the biosynthesis of glycopeptides, beta-lactams, and aminoglycosides, among others, concurrent with the high abundance of ARGs against said antibiotics. Additionally, we identified BGCs with a predicted antibacterial function in both eukaryotes and bacteria. While a limited number of studies such as Waschulin *et al*. [54] and Liao *et al*. [55], have shown BGCs in pristine environments, none of these studies have contextualized the co-occurrence of BGCs with AMR. Hence, we not only found that most of our MAGs contain BGCs, of which many have an antibacterial function, but also found all MAGs to encode multiple resistance genes. Additionally, we found several BGCs closely localized to ARGs on the same contig, thereby indicating an immediate self-defense mechanism against the encoded secondary metabolites. This agrees with the resistance hypothesis highlighted by Tran *et al*. stating that a gene conferring resistance to potentially harmful metabolites produced by the organism are to be found within the BGC-encoding operons [14]. We also observed that the recently identified CPR bacteria [56] (in our case, phylum Patescibacteria) not only encoded for AMR but also harboured genes associated with the production of molecules with antibacterial effects. Although Patescibacteria have been identified in oligotrophic environments [57,58] with carbon and/or nutrient limitations similar to those observed for GFSs, it is plausible that their ability to survive with minimal biosynthetic and metabolic pathways may indeed depend on the expression of BGCs and AMR. At the time of writing, a preprint by Maatouk *et al*. [59], described the presence of ARGs across publicly available CPR bacterial genomes. In addition, we report the identification of AMR within GFS-derived CPR genomes, likely as a means of competitive inhibition against other taxa. Alternatively, biofilms may also allow for collective resistance, tolerance, and exposure protection to antibacterial compounds [60]. The AMR and BGCs encoded by most phyla may therefore affect cooperation and/or interactions associated with nutrient exchange, leading to the privatization of public goods [60]. Such a phenomenon may be achieved due to the competition within taxa, both at the intra- and inter-species levels, via secretion of toxins [53] and occupying spatial niches [61,62] thereafter. Furthermore, Stubbendieck and Straight previously highlighted the multifaceted effects of bacterial competition which include the potential taxation and subsequent increase in bacterial fitness [63]. Thus, the *in-situ* competition within multi-species biofilms may allow for cross-phyla and cross-domain interactions whilst simultaneously increasing the overall fitness of the endogenous epilithic microbial community. Alternatively, these interactions or lack thereof may shape the overall community including spatial organisation [64], especially in energy limited systems such as the GFSs.

## Conclusions

Epilithic biofilms are an integral and key mode of survival in extreme environments such as glacier-fed stream ecosystems. Herein, we report that these biofilms provide critical insights into the naturally occurring resistome. Our findings demonstrate that intra- and inter-domain competition and survival mechanisms shed light on the ecological dimension of microbial communities. Furthermore, we reveal the congruence of genes encoding for both BGCs and AMR, in both bacteria and eukaryotes. More importantly, we highlight for the first time the comprehensive AMR profile of CPR bacteria and of (micro-)eukaryotes. Collectively, our results highlight underlying resistance mechanisms, including BGCs, employed in ‘biological warfare’ in oligotrophic and challenging glacier-fed stream ecosystems.

## List of Abbreviations

AMR: Antimicrobial resistance
ARGs: Antimicrobial resistance gene(s)
BGC: Biosynthetic gene clusters
CA: Caucasus
CPR: Candidate Phyla radiation
GFSs: Glacier-fed stream(s)
GL: Glacier
IRS-RS: isoleucyl-tRNA synthetase - high resistance
IMP: Integrate Meta-Omics Pipeline
KEGG: Kyoto Encyclopedia of Genes and Genomes
MAGs: Metagenome-assembled genome(s)
NRPS: Non-ribosomal peptide synthetases
PKS: Polyketide synthases (type I and type II)
RiPPs: Post-translationally modified peptide(s)
SA: Southern Alps

## Declarations

Ethics approval and consent to participate

Not applicable

Consent for publication

Not applicable

## Availability of data and material

The Biosample accession IDs listed under Supp. Table 3 can be found on NCBI under the BioProject accession# **PRJNA733707**. The analyses code for IMP and downstream analyses is detailed at https://git-r3lab.uni.lu/susheel.busi/nomis_pipeline. Binning and manual refinement of eukaryotic MAGs was done as described here: https://git-r3lab.uni.lu/susheel.busi/nomis_pipeline/-/blob/master/workflow/notes/MiscEUKMAGs.md. All visualization and analysis code is available at: https://git-r3lab.uni.lu/laura.denies/Rock_Biofilm_AMR.

## Authors’ contributions

SBB, LdN, PW, and TJB conceived the project. PP extracted DNA, SBB and PP prepared the metagenomic libraries for sequencing. SBB and LdN conceptualized and performed the data analyses. SBB and LdN wrote the manuscript with PW and TJB, with significant input and editing from all coauthors.

## Acknowledgements

We gratefully acknowledge the laboratory support from Emmy Marie Oppliger at EPFL and Lea Grandmougin, Janine Habier, Laura Lebrun at the University of Luxembourg. We also acknowledge the key input from Rashi Halder at the LCSB Sequencing Platform regarding library preparation. We thank Patrick May and Cedric Christian Laczny for the crucial insights into metagenomic processing. The computational analyses were performed at the HPC facilities at the University of Luxembourg (https://hpc.uni.lu) [65].

---

**Supplementary figure 1.**
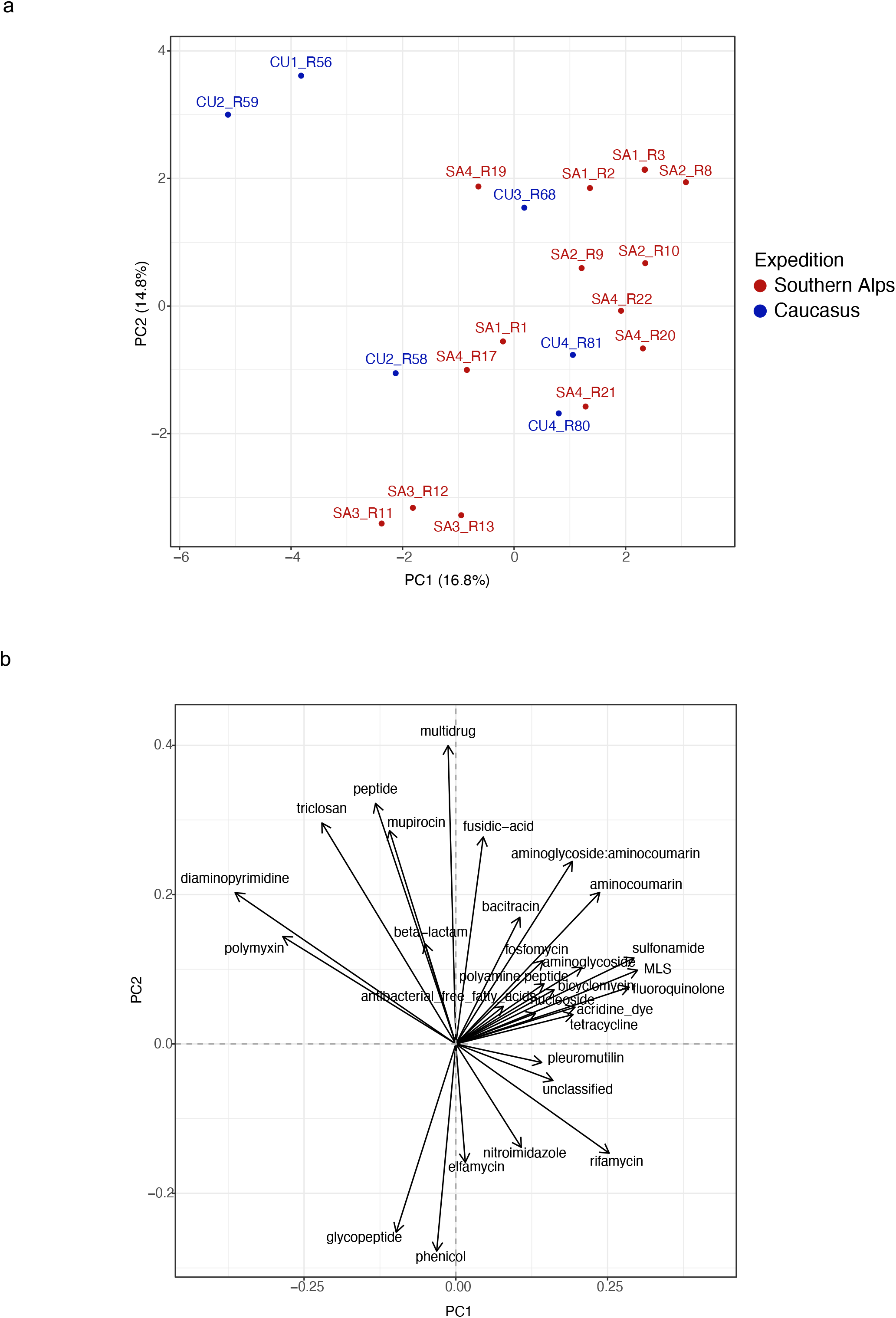
Ordination analyses reveal the (dis)similarity of the GFS resistomes. (a) Principal component analyses depicting the overall similarity of the individual GFS resistomes. Each dot represents the resistome predicted from a single metagenome. SA: Southern Alps. CU: Caucasus. (b) Biplot demonstrating the underlying factors, i.e. ARG abundances across 29 AMR categories, driving the similarity within the GFS epilithic resistomes.

**Supplementary figure 2.**
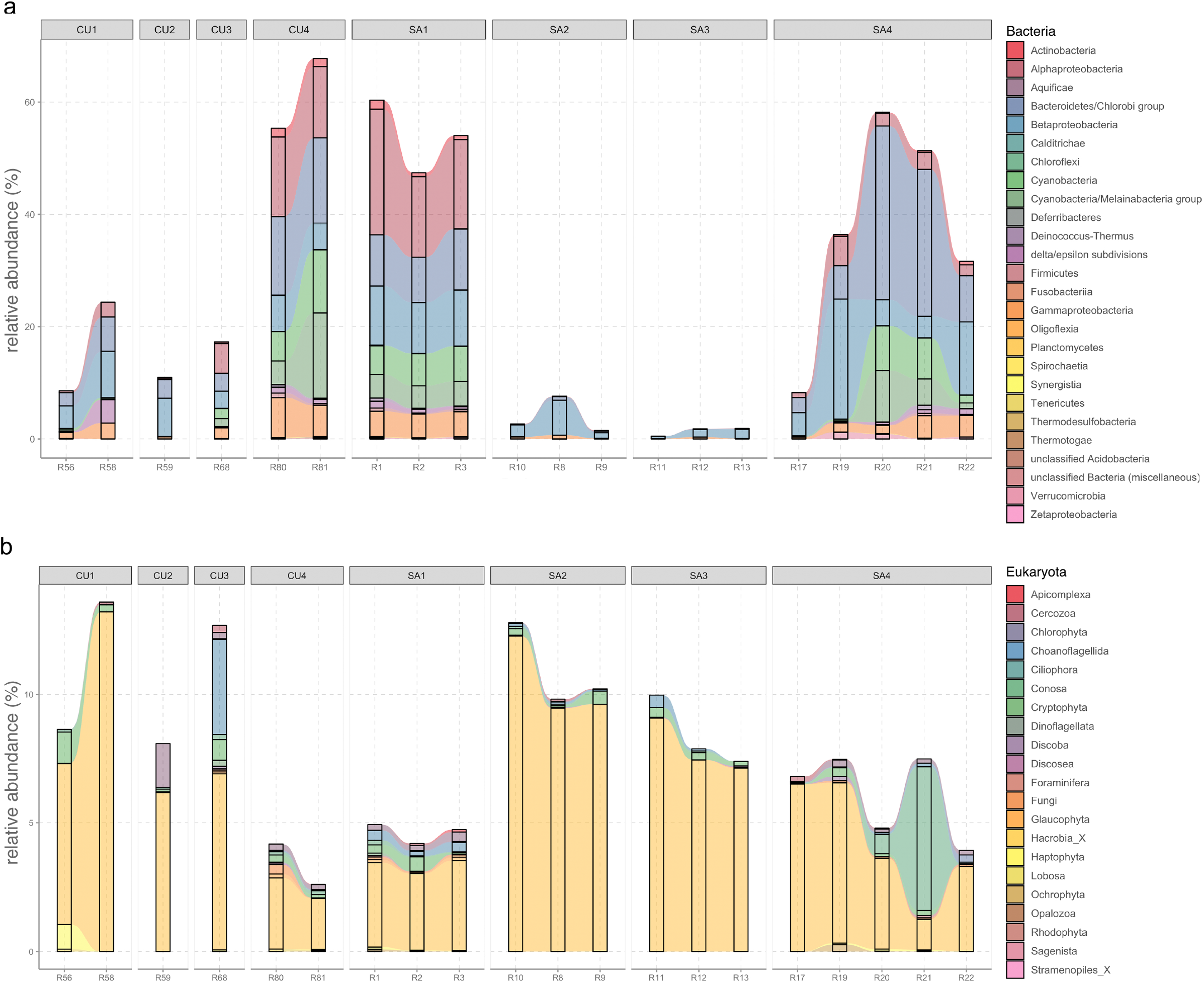
Bacterial and eukaryotic phyla encode AMR. (a) Relative abundance of the bacteria associated with AMR. The stacked bar plots are facetted by the individual GFSs where the epilithic biofilms were collected. The colors represent the individual phyla. (b) Stacked bar plots indicating the relative abundance of the AMR encoded by eukaryotes.

**Supplementary figure 3.**
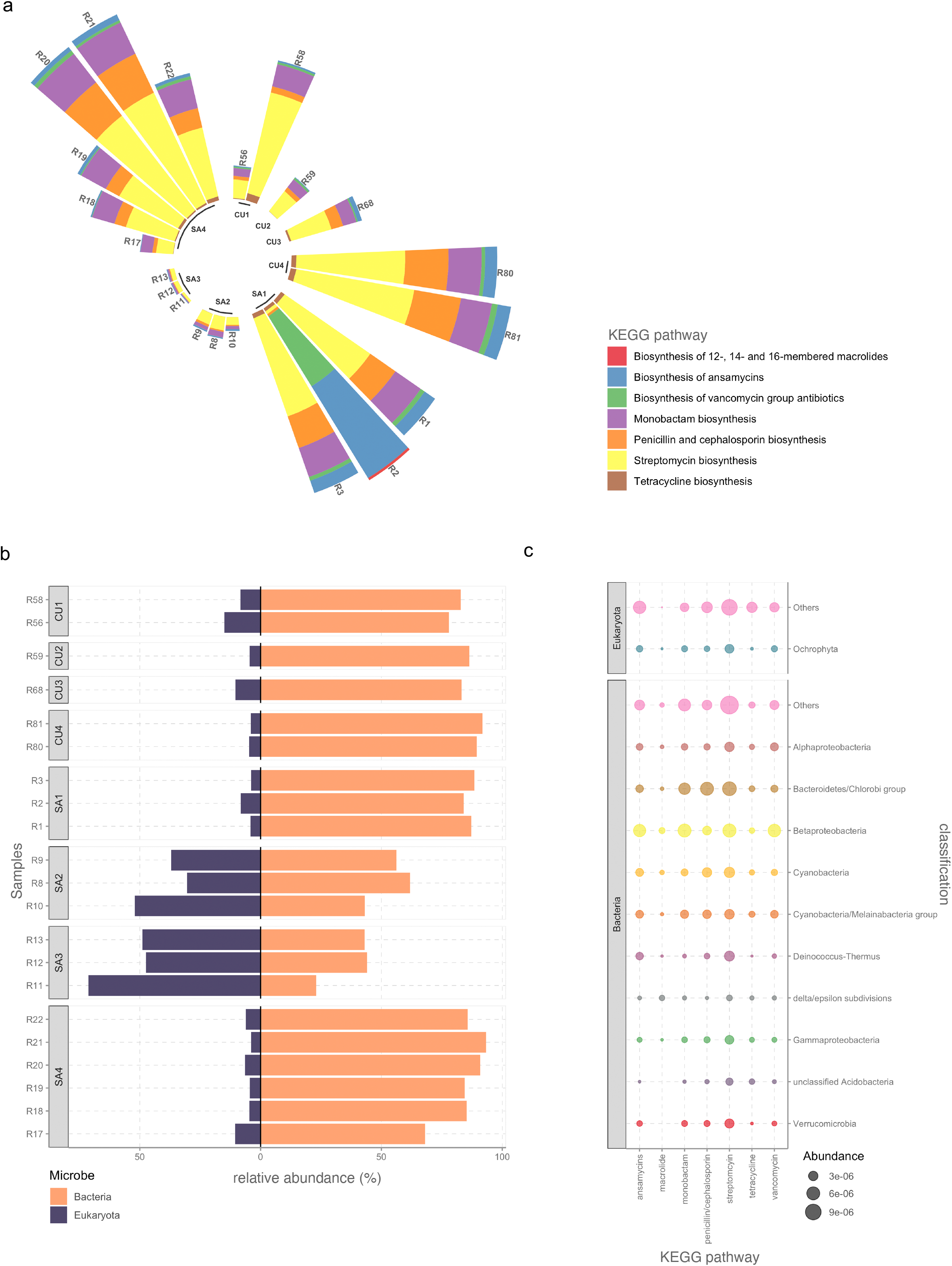
Antibiotic synthesis pathway assessment via KEGG orthology. (a) Relative abundance of KEGG pathways associated with antibiotic synthesis across the 21 epilithic biofilms. (b) Bar plots indicating the relative abundance of the antibiotic associated KEGG pathways mediated by bacteria and eukaryotes. (c) Normalised relative abundance of pathways associated with antibiotic production in the KEGG database, juxtaposed with the various phyla encoding these genes.

## Supplementary data

**Supplementary table 1. Sample metadata**

**Supplementary table 2. List of ARGs identified across 21 GFS epilithic biofilms**

**Supplementary table 3. NCBI accession metadata**

## References

1. Stanton IC, Bethel A, Leonard AFC, Gaze WH, Garside R. What is the research evidence for antibiotic resistance exposure and transmission to humans from the environment? A systematic map protocol. Environ Evid. 2020;9: 12.

2. Balasegaram M. Learning from COVID-19 to Tackle Antibiotic Resistance. ACS Infect Dis. 2021;7: 693–694.

3. Wright GD. The antibiotic resistome: the nexus of chemical and genetic diversity. Nat Rev Microbiol. 2007;5: 175–186.

4. D’Costa VM, King CE, Kalan L, Morar M, Sung WWL, Schwarz C, et al. Antibiotic resistance is ancient. Nature. 2011;477: 457–461.

5. Scott LC, Lee N, Aw TG. Antibiotic Resistance in Minimally Human-Impacted Environments. Int J Environ Res Public Health. 2020;17. doi:10.3390/ijerph17113939

6. Tyc O, Song C, Dickschat JS, Vos M, Garbeva P. The Ecological Role of Volatile and Soluble Secondary Metabolites Produced by Soil Bacteria. Trends Microbiol. 2017;25: 280–292.

7. Chen R, Wong HL, Kindler GS, MacLeod FI, Benaud N, Ferrari BC, et al. Discovery of an Abundance of Biosynthetic Gene Clusters in Shark Bay Microbial Mats. Front Microbiol. 2020;11: 1950.

8. Demain AL, Fang A. The Natural Functions of Secondary Metabolites. In: Fiechter A, editor. History of Modern Biotechnology I. Berlin, Heidelberg: Springer Berlin Heidelberg; 2000. pp. 1–39.

9. Newman DJ, Cragg GM. Natural Products as Sources of New Drugs from 1981 to 2014. J Nat Prod. 2016;79: 629–661.

10. Medema MH, Kottmann R, Yilmaz P, Cummings M, Biggins JB, Blin K, et al. Minimum Information about a Biosynthetic Gene cluster. Nat Chem Biol. 2015;11: 625–631.

11. Martinet L, Naômé A, Deflandre B, Maciejewska M, Tellatin D, Tenconi E, et al. A Single Biosynthetic Gene Cluster Is Responsible for the Production of Bagremycin Antibiotics and Ferroverdin Iron Chelators. MBio. 2019;10. doi:10.1128/mBio.01230-19

12. Martínez-Núñez MA, López VEL y. Nonribosomal peptides synthetases and their applications in industry. Sustainable Chemical Processes. 2016;4: 1–8.

13. Ridley CP, Lee HY, Khosla C. Evolution of polyketide synthases in bacteria. Proc Natl Acad Sci U S A. 2008;105: 4595–4600.

14. Tran PN, Yen M-R, Chiang C-Y, Lin H-C, Chen P-Y. Detecting and prioritizing biosynthetic gene clusters for bioactive compounds in bacteria and fungi. Appl Microbiol Biotechnol. 2019;103: 3277–3287.

15. Cundliffe E, Bate N, Butler A, Fish S, Gandecha A, Merson-Davies L. The tylosin-biosynthetic genes of Streptomyces fradiae. Antonie Van Leeuwenhoek. 2001;79: 229–234.

16. Kwun MJ, Hong H-J. Genome Sequence of Streptomyces toyocaensis NRRL 15009, Producer of the Glycopeptide Antibiotic A47934. Genome Announc. 2014;2. doi:10.1128/genomeA.00749-14

17. Busi SB, Bourquin M, Fodelianakis S, Michoud G, Kohler TJ, Peter H, et al. Genomic and metabolic adaptations of biofilms to ecological windows of opportunities in glacier-fed streams. bioRxiv. 2021. p. 2021.10.07.463499. doi:10.1101/2021.10.07.463499

18. Battin TJ, Besemer K, Bengtsson MM, Romani AM, Packmann AI. The ecology and biogeochemistry of stream biofilms. Nat Rev Microbiol. 2016;14: 251–263.

19. Battin TJ, Wille A, Sattler B, Psenner R. Phylogenetic and functional heterogeneity of sediment biofilms along environmental gradients in a glacial stream. Appl Environ Microbiol. 2001;67: 799–807.

20. Gaynes R. The Discovery of Penicillin—New Insights After More Than 75 Years of Clinical Use. Emerg Infect Dis. 2017;23: 849.

21. Netzker T, Flak M, Krespach MK, Stroe MC, Weber J, Schroeckh V, et al. Microbial interactions trigger the production of antibiotics. Curr Opin Microbiol. 2018;45: 117–123.

22. Busi SB, Pramateftaki P, Brandani J, Fodelianakis S, Peter H, Halder R, et al. Optimised biomolecular extraction for metagenomic analysis of microbial biofilms from high-mountain streams. PeerJ. 2020;8: e9973.

23. Narayanasamy S, Jarosz Y, Muller EEL, Heintz-Buschart A, Herold M, Kaysen A, et al. IMP: a pipeline for reproducible reference-independent integrated metagenomic and metatranscriptomic analyses. Genome Biol. 2016;17: 260.

24. de Nies L, Lopes S, Busi SB, Galata V, Heintz-Buschart A, Laczny CC, et al. PathoFact: a pipeline for the prediction of virulence factors and antimicrobial resistance genes in metagenomic data. Microbiome. 2021;9: 49.

25. Alcock BP, Raphenya AR, Lau TTY, Tsang KK, Bouchard M, Edalatmand A, et al. CARD 2020: antibiotic resistome surveillance with the comprehensive antibiotic resistance database. Nucleic Acids Res. 2020;48: D517–D525.

26. Liao Y, Smyth GK, Shi W. featureCounts: An efficient general-purpose program for assigning sequence reads to genomic features. arXiv [q-bio.GN]. 2013. Available: http://arxiv.org/abs/1305.3347

27. Yoon B-J. Hidden Markov Models and their Applications in Biological Sequence Analysis. Curr Genomics. 2009;10: 402–415.

28. Eddy SR. Accelerated Profile HMM Searches. PLoS Comput Biol. 2011;7: e1002195.

29. Blin K, Shaw S, Kloosterman AM, Charlop-Powers Z, van Wezel GP, Medema MH, et al. antiSMASH 6.0: improving cluster detection and comparison capabilities. Nucleic Acids Res. 2021;49: W29–W35.

30. Hannigan GD, Prihoda D, Palicka A, Soukup J, Klempir O, Rampula L, et al. A deep learning genome-mining strategy for biosynthetic gene cluster prediction. Nucleic Acids Res. 2019;47: e110.

31. Hu Y, Yang X, Qin J, Lu N, Cheng G, Wu N, et al. Metagenome-wide analysis of antibiotic resistance genes in a large cohort of human gut microbiota. Nat Commun. 2013;4: 2151.

32. Computing R, Others. R: A language and environment for statistical computing. Vienna: R Core Team. 2013. Available: https://www.yumpu.com/en/document/view/6853895/r-a-language-and-environment-for-statistical-computing

33. Wickham H, Averick M, Bryan J, Chang W, McGowan L, François R, et al. Welcome to the tidyverse. J Open Source Softw. 2019;4: 1686.

34. Brunson J. ggalluvial: Layered Grammar for Alluvial Plots. J Open Source Softw. 2020;5: 2017.

35. Gu Z, Eils R, Schlesner M. Complex heatmaps reveal patterns and correlations in multidimensional genomic data. Bioinformatics. 2016;32: 2847–2849.

36. Krause KM, Serio AW, Kane TR, Connolly LE. Aminoglycosides: An Overview. Cold Spring Harb Perspect Med. 2016;6. doi:10.1101/cshperspect.a027029

37. Tahlan K, Jensen SE. Origins of the β-lactam rings in natural products. J Antibiot. 2013;66: 401–410.

38. Borges-Walmsley MI, McKeegan KS, Walmsley AR. Structure and function of efflux pumps that confer resistance to drugs. Biochem J. 2003;376: 313–338.

39. Kanehisa M, Goto S. KEGG: Kyoto Encyclopedia of Genes and Genomes. Nucleic Acids Res. 2000;28: 27–30.

40. Tortorella E, Tedesco P, Palma Esposito F, January GG, Fani R, Jaspars M, et al. Antibiotics from Deep-Sea Microorganisms: Current Discoveries and Perspectives. Mar Drugs. 2018;16. doi:10.3390/md16100355

41. McCann CM, Christgen B, Roberts JA, Su J-Q, Arnold KE, Gray ND, et al. Understanding drivers of antibiotic resistance genes in High Arctic soil ecosystems. Environ Int. 2019;125: 497–504.

42. Yuan K, Yu K, Yang R, Zhang Q, Yang Y, Chen E, et al. Metagenomic characterization of antibiotic resistance genes in Antarctic soils. Ecotoxicol Environ Saf. 2019;176: 300–308.

43. Centurion VB, Delforno TP, Lacerda-Júnior GV, Duarte AWF, Silva LJ, Bellini GB, et al. Unveiling resistome profiles in the sediments of an Antarctic volcanic island. Environ Pollut. 2019;255: 113240.

44. Van Goethem MW, Pierneef R, Bezuidt OKI, Van De Peer Y, Cowan DA, Makhalanyane TP. A reservoir of “historical” antibiotic resistance genes in remote pristine Antarctic soils. Microbiome. 2018;6: 40.

45. Brown JR, Zhang J, Hodgson JE. A bacterial antibiotic resistance gene with eukaryotic origins. Curr Biol. 1998;8: R365–7.

46. Fairlamb AH, Gow NAR, Matthews KR, Waters AP. Drug resistance in eukaryotic microorganisms. Nat Microbiol. 2016;1: 16092.

47. Silva A, Silva SA, Carpena M, Garcia-Oliveira P, Gullón P, Barroso MF, et al. Macroalgae as a Source of Valuable Antimicrobial Compounds: Extraction and Applications. Antibiotics (Basel). 2020;9. doi:10.3390/antibiotics9100642

48. Martins RM, Nedel F, Guimarães VBS, da Silva AF, Colepicolo P, de Pereira CMP, et al. Macroalgae Extracts From Antarctica Have Antimicrobial and Anticancer Potential. Front Microbiol. 2018;9: 412.

49. Karkman A, Pärnänen K, Larsson DGJ. Fecal pollution can explain antibiotic resistance gene abundances in anthropogenically impacted environments. Nat Commun. 2019;10: 80.

50. Antelo V, Giménez M, Azziz G, Valdespino-Castillo P, Falcón LI, Ruberto LAM, et al. Metagenomic strategies identify diverse integron-integrase and antibiotic resistance genes in the Antarctic environment. Microbiologyopen. 2021;10. doi:10.1002/mbo3.1219

51. Hernández F, Calisto-Ulloa N, Gómez-Fuentes C, Gómez M, Ferrer J, González-Rocha G, et al. Occurrence of antibiotics and bacterial resistance in wastewater and sea water from the Antarctic. J Hazard Mater. 2019;363: 447–456.

52. Reygaert WC. An overview of the antimicrobial resistance mechanisms of bacteria. AIMS Microbiol. 2018;4: 482–501.

53. Granato ET, Meiller-Legrand TA, Foster KR. The Evolution and Ecology of Bacterial Warfare. Curr Biol. 2019;29: R521–R537.

54. Waschulin V, Borsetto C, James R, Newsham KK, Donadio S, Corre C, et al. Biosynthetic potential of uncultured Antarctic soil bacteria revealed through long-read metagenomic sequencing. ISME J. 2021. doi:10.1038/s41396-021-01052-3

55. Liao L, Su S, Zhao B, Fan C, Zhang J, Li H, et al. Biosynthetic Potential of a Novel Antarctic Actinobacterium Marisediminicola antarctica ZS314T Revealed by Genomic Data Mining and Pigment Characterization. Mar Drugs. 2019;17. doi:10.3390/md17070388

56. Hug LA, Baker BJ, Anantharaman K, Brown CT, Probst AJ, Castelle CJ, et al. A new view of the tree of life. Nat Microbiol. 2016;1: 16048.

57. Tian R, Ning D, He Z, Zhang P, Spencer SJ, Gao S, et al. Small and mighty: adaptation of superphylum Patescibacteria to groundwater environment drives their genome simplicity. Microbiome. 2020;8: 51.

58. Vigneron A, Cruaud P, Langlois V, Lovejoy C, Culley AI, Vincent WF. Ultra-small and abundant: Candidate phyla radiation bacteria are potential catalysts of carbon transformation in a thermokarst lake ecosystem. Limnol Oceanogr Lett. 2020;5: 212–220.

59. Maatouk M, Ibrahim A, Rolain J-M, Merhej V, Bittar F. Small and equipped: the rich repertoire of antibiotic resistance genes in Candidate Phyla Radiation genomes. bioRxiv. 2021. p. 2021.07.02.450847. doi:10.1101/2021.07.02.450847

60. Bottery MJ, Pitchford JW, Friman V-P. Ecology and evolution of antimicrobial resistance in bacterial communities. ISME J. 2021;15: 939–948.

61. Bottery MJ, Passaris I, Dytham C, Wood AJ, van der Woude MW. Spatial Organization of Expanding Bacterial Colonies Is Affected by Contact-Dependent Growth Inhibition. Curr Biol. 2019;29: 3622–3634.e5.

62. Schluter J, Nadell CD, Bassler BL, Foster KR. Adhesion as a weapon in microbial competition. ISME J. 2015;9: 139–149.

63. Stubbendieck RM, Straight PD. Multifaceted Interfaces of Bacterial Competition. J Bacteriol. 2016;198: 2145–2155.

64. Estrela S, Brown SP. Community interactions and spatial structure shape selection on antibiotic resistant lineages. PLoS Comput Biol. 2018;14: e1006179.

65. Varrette S, Bouvry P, Cartiaux H, Georgatos F. Management of an academic HPC cluster: The UL experience. 2014 International Conference on High Performance Computing Simulation (HPCS). 2014. pp. 959–967.

